# Flagellar structures from the bacterium *Caulobacter crescentus* and implications for phage ϕCbK predation of multi-flagellin bacteria

**DOI:** 10.1101/2020.07.27.223248

**Authors:** Eric J. Montemayor, Nicoleta T. Ploscariu, Juan C. Sanchez, Daniel Parrell, Rebecca S. Dillard, Conrad W. Shebelut, Zunlong Ke, Ricardo C. Guerrero-Ferreira, Elizabeth R. Wright

**Affiliations:** Department of Biochemistry, University of Wisconsin, Madison, WI 53706; Cryo-Electron Microscopy Research Center, Department of Biochemistry, University of Wisconsin, Madison, WI 53706; Biophysics Graduate Program, University of Wisconsin, Madison, WI 53706; Division of Infectious Diseases, Department of Pediatrics, Emory University School of Medicine, Children’s Healthcare of Atlanta, Atlanta, GA, 30322; Department of Pathology and Laboratory Medicine, Emory University School of Medicine, Atlanta, GA, 30322; School of Biological Sciences, Georgia Institute of Technology, Atlanta, GA, 30332; Morgridge Institute for Research, Madison, WI, 53715

**Author notes:** These authors contributed equally (Juan C. Sanchez, Daniel Parrell).

## Abstract

*Caulobacter crescentus* is a gram-negative alpha-proteobacterium that commonly lives in oligotrophic fresh and salt-water environments. *C. crescentus* is a host to many bacteriophages, including ϕCbK and ϕCbK-like bacteriophages, which first adsorb to cells by interaction with the bacterial flagellum. It is commonly thought that the six paralogs of the flagellin gene present in *C. crescentus* are important for bacteriophage evasion. Here, we show that deletion of specific flagellins in *C. crescentus* can indeed attenuate ϕCbK adsorption efficiency, although no single deletion completely ablates ϕCbK adsorption. Thus, bacteriophage ϕCbK likely recognizes a common motif amongst the six known flagellins in *C. crescentus* with varying degrees of efficiency. Interestingly, we observe that most deletion strains still generate flagellar filaments, with the exception of a strain that contains only the most divergent flagellin, FljJ, or a strain that contains only FljN and FljO. To visualize the surface residues that are likely recognized by ϕCbK, we determined two high-resolution structures of the FljK filament, with and without an amino acid substitution that induces straightening of the filament. We observe post-translational modifications on conserved surface threonine residues of FljK that are likely O-linked glycans. The possibility of interplay between these modifications and ϕCbK adsorption is discussed. We also determined the structure of a filament composed of a heterogeneous mixture of FljK and FljL, the final resolution of which was limited to approximately 4.6 Å. Altogether, this work builds a platform for future investigation of how phage ϕCbK infects *C. crescentus* at the molecular level.

## INTRODUCTION

Bacterial motility relies upon signaling cascades that link chemical signals to mechanical responses in various bacterial appendages. One of the most dynamic appendages is the bacterial flagellum, which is assembled from over sixty structural and regulatory genes (1, 2). The flagellum is a multiprotein complex with three distinct regions: the motor, hook, and filament (2, 3). The motor, also called the basal body, is located within the cell wall and generates torque via an electrochemical potential gradient, resulting in motor speeds of between 300 and 1700 Hz (4–6). This mechanical signal is transferred to the filament by the flagella’s universal joint called the hook, a tubular supercoiled helical structure (7). Motility is achieved by the flagellar filament, which acts as an Archimedes’ screw to propel the cell body (8).

The flagellar filament is a long cylindrical structure with a diameter of 12-15 nm that is composed of thousands of flagellin subunits, and as such can span approximately 15 μm in length (9). Individual flagellin subunits typically have masses between 25 kDa and 65 kDa and consist of multiple domains (10, 11). The two most conserved domains, denoted D0 and D1, are associated with flagellin polymerization and form the lumen of the flagellar filament. Specific regions of D1 are known recognition sites for the innate immune systems of mammals and plants (12–15). However, the two surface exposed domains, D2 and D3, demonstrate high sequence variation (15, 16) and are known loci for a myriad of post-translational modifications, such as O- and N-linked glycosylation (17–20). Mass spectroscopy studies of flagellar proteins show that *Halobacterium* and *Clostridium* flagellins are among the most heavily glycosylated species, where glycans account for 10% of their molecular weight (21). It was thought that only O-linked glycans were found in bacteria, but studies show that both O-linked and N-linked glycans are present in different bacterial species, similar to the homologous archaeal flagella (22–24). The D2 and D3 regions have also been shown to serve as an initial attachment point between flagellotropic bacteriophages and their hosts (25–30). Thus, the periphery of flagellins serves as a hub for post-translational modifications and protein-protein interactions that mediate bacteriophage predation.

The process of bacteriophage adsorption, where a bacteriophage attaches to and infects a cell, is dependent upon specific surface interactions and physical diffusion (Fig. 1A,B) (31). In most cases, phage tail fibers wrap directly around the flagellum as the bacteria swim (32–35). Previously, we characterized an alternative, head-first, attachment mechanism of the bacteriophage ϕCbK in *C. crescentus* (30). In this mechanism, the phage head filament wraps around the flagellum allowing for the localization of bacteriophages to the cell pole as the flagellar filament rotates (30). Next, the tail of ϕCbK binds to the pilus filament and then is retracted to the pilus portal for irreversible attachment (30). However, much remains unknown regarding the underlying adhesion mechanism and the structural contacts that endow specificity between phage ϕCbK and the *C. crescentus* flagellum. For example, which residues on the surface of the flagellum and individual flagellins directly interact with the ϕCbK head filament? Answering this question has been impeded by the lack of a high-resolution model of the *C. crescentus* flagellar filament.

**Figure 1.**
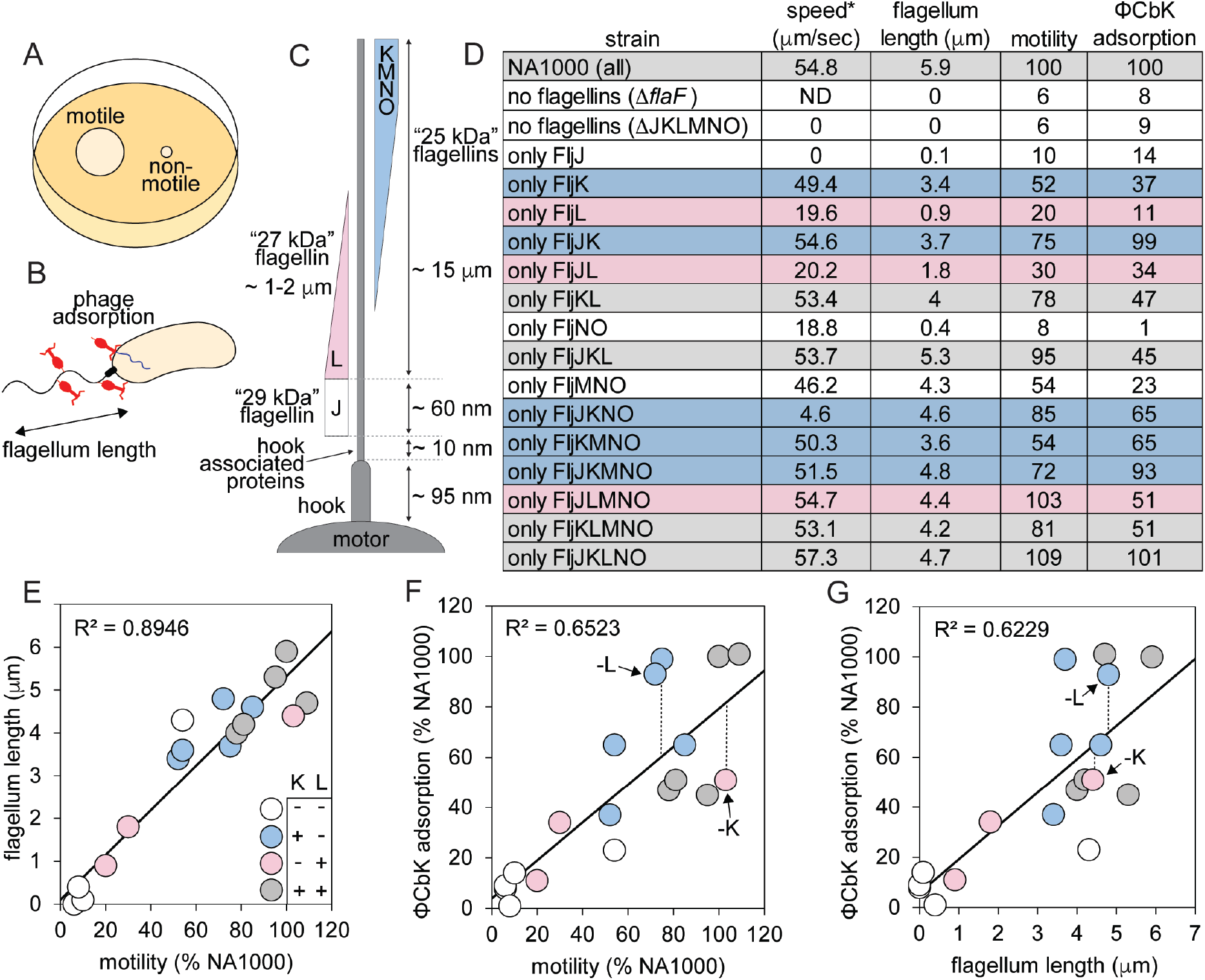
Contribution of various flagellins to *C. crescentus* mobility and phage adsorption. A) The motility assay used here tracks colony growth across a low percent agar medium. Non-motile colonies spread less than motile. B) Schematic of phage adsorption to *C. crescentus*, which includes attachment to the flagella, subsequent attachment to the cell, and genome ejection. The length of flagella in this work was determined by negative stain electron microscopy of *C. crescentus*. C) Schematic of known organization of flagellin proteins in the *C. crescentus* flagellum. D) Summary of functional assays performed here. For clarity, *C. crescentus* strains are designated by the flagellin(s) they synthesize. The data for cell swim speed are reproduced from a previous work for comparison and are denoted by an asterisk (37). Data for motility and adsorption are normalized to that of wild type *C. crescentus* strain NA1000. Blue strains contain FljK but not FljL. Pink strains contain FljL but not FljK. Grey and white strains contain both or neither of FljK and FljL, respectively. E-G) Plots showing correlation between observed phenotypes. Strains are colored as in panel D.

The number of flagellin genes present in bacterial genomes is highly variable. For example, some bacteria have a single flagellin gene, such as *Escherichia coli* and *Bacillus subtilis*, while others maintain multiple flagellin genes, including *Helicobacter pylori, Rhizobium meliloti, Salmonella enterica* serovar *Typhimurium*, and the *Vibrionaceae* (36). In the case of *S. enterica* serovar *Typhimurium*, phase variation is used to control which flagellin is incorporated into flagellar filaments, and as such single filaments contain only one type of flagellin protein. However, almost 50% of bacterial species, including *C. crescentus*, assemble single flagellar filaments that are compositionally heterogeneous (37). The functional purpose of this heterogeneity is unknown, although it may provide a selective advantage for responding to environmental pressures, improved pathogenicity, or resistance to bacteriophage predation. The latter has already been demonstrated in bacteria that exhibit phase variation (38–42).

The *C. crescentus* flagellum has been a model for understanding the composition and structure of bacterial flagella (43–45). The *C. crescentus* flagellar filament is composed of six independent flagellin subunits (46). Immuno-EM labeling of such filaments suggests that individual subunits are differentially incorporated along the length of the filament (47–49) (Fig. 1C). The flagellins are expressed from two separate loci on the *C. crescentus* genome. The alpha locus contains the *fljJ* (29 kDa flagellin), *fljK* (25 kDa flagellin) and *fljL* (27 kDa flagellin) genes as well as the two flagellin translation regulators, *flaF* and *flbT*. The beta locus includes the *fljM, fljN*, and *fljO* genes, all of which encode 25 kDa flagellins (37, 46). Previously, it was determined that alterations to the six flagellins incorporated into the *C. crescentus* flagellum could reduce but not enhance motility (37).

The small size of flagellin monomers and high degree of symmetry in assembled filaments provides challenges for high-resolution structure determination by contemporary cryo-electron microscopy and helical reconstruction methods (50–52).

Typically, it is necessary to determine an initial estimate of helical symmetry by *de novo* indexing. However, this requires long and straight filaments, or 2-D classes of smaller “boxes” with sufficient signal in their amplitude spectra for layer line indexing. When it is not possible to index a helical assembly *de novo*, initial estimates of helical symmetry can be obtained from homologous structures (50).

The architecture of flagella are well conserved across bacterial species. Most contain 11-start helices with protofilaments that run nearly parallel to the helical axis, with the exception of *C. jejuni* which contains a 7-start helix (53–55). A handful of 11-start flagellin structures have been determined, including flagellar filaments from *B. subtilis, S. enterica, Pseudomonas aeruginosa*, and *Kurthia* sp. strain 11kri321 (9, 54–56). In the case of the *B. subtilis, S. enterica*, and *P. aeruginosa* reconstructions, single residue substitutions were used to straighten the filaments by locking the 11-start protofilament into either right (R-type) or left (L-type) handedness conformations, which yielded a uniform filament onto which helical symmetry could be more effectively determined and applied during structure determination (55, 56). In *Kurthia*, straightening substitutions were not employed to intentionally lock the filament into a particular R-type or L-type handedness (54).

We have previously demonstrated that reduced motility of *C. crescentus* leads to decreased adsorption of phage ϕCbK (4, 30), as similarly observed with *E. coli* phage x (2, 28), *B. subtilis* phage PBS1 (56, 57), and *Agrobacterium tumefaciens* phages GS2 and GS6 (28, 58). Here we characterize the effects of flagellin composition in *C. crescentus* on phage ϕCbK adsorption. These results were achieved by combining motility and bacteriophage adsorption assays with negative stain transmission electron microscopy (TEM), cryo-electron microscopy (cryo-EM), and cryo-electron tomography (cryo-ET) of *C. crescentus* mutants. We show that no one flagellin protein is essential for phage adsorption, although a preference for some flagellins is apparent. As such, bacteriophage ϕCbK likely recognizes a common motif amongst the six *C. crescentus* flagellins. We also present the first high-resolution helical reconstructions for the FljK flagellar filament from *C. crescentus* and a lower-resolution model of a flagellar filament containing more than one flagellin.

## RESULTS

Since *C. crescentus* harbors six distinct flagellins, we sought to deconvolute the effect of motility, filament composition, and structure on ϕCbK adsorption, by using a series of *C. crescentus* mutant strains that lacked one, several, or all but one flagellin (Fig. 1D). For clarity, strains will be referred to by the flagellin(s) that they synthesize.

### Phage adsorption to flagellin mutants of *C. crescentus*

*C. crescentus* strains lacking flagellar filaments altogether *(ΔflaF* and *ΔfljJKLMNO)* exhibit reduced motility and adsorption rates of ϕCbK (Fig. 1D). We also find that motility and filament length are strongly correlated. These two results, which agree with previous observations (30, 37, 59), serve as an internal control for the analyses conducted here (Fig. 1E). Interestingly, we observe weaker correlations between motility and adsorption, and also between filament length and adsorption, suggesting motility or filament length alone do not exclusively drive adsorption (Fig. 1F,G), and as such, phage ϕCbK is likely sensitive to composition of the flagellar filament.

In all strains tested, expression of a single flagellin protein reduced ϕCbK adsorption. Adsorption to FljJ and FljL, was extremely low at ~10% of wild type (Figs. 1D, S1 and S2). This reduction was accompanied by reduced motility and swimming speed (37), as well as relatively short flagellum length (Fig. 2). In the case of FljK, ϕCbK adsorption was reduced to ~35% and motility of the strain was ~50% of wild type, although swimming speed (37) and flagellum length remained similar to that of wild type. Interestingly, the FljJ strain lacked filaments altogether (Fig. 2B).

**Figure 2.**
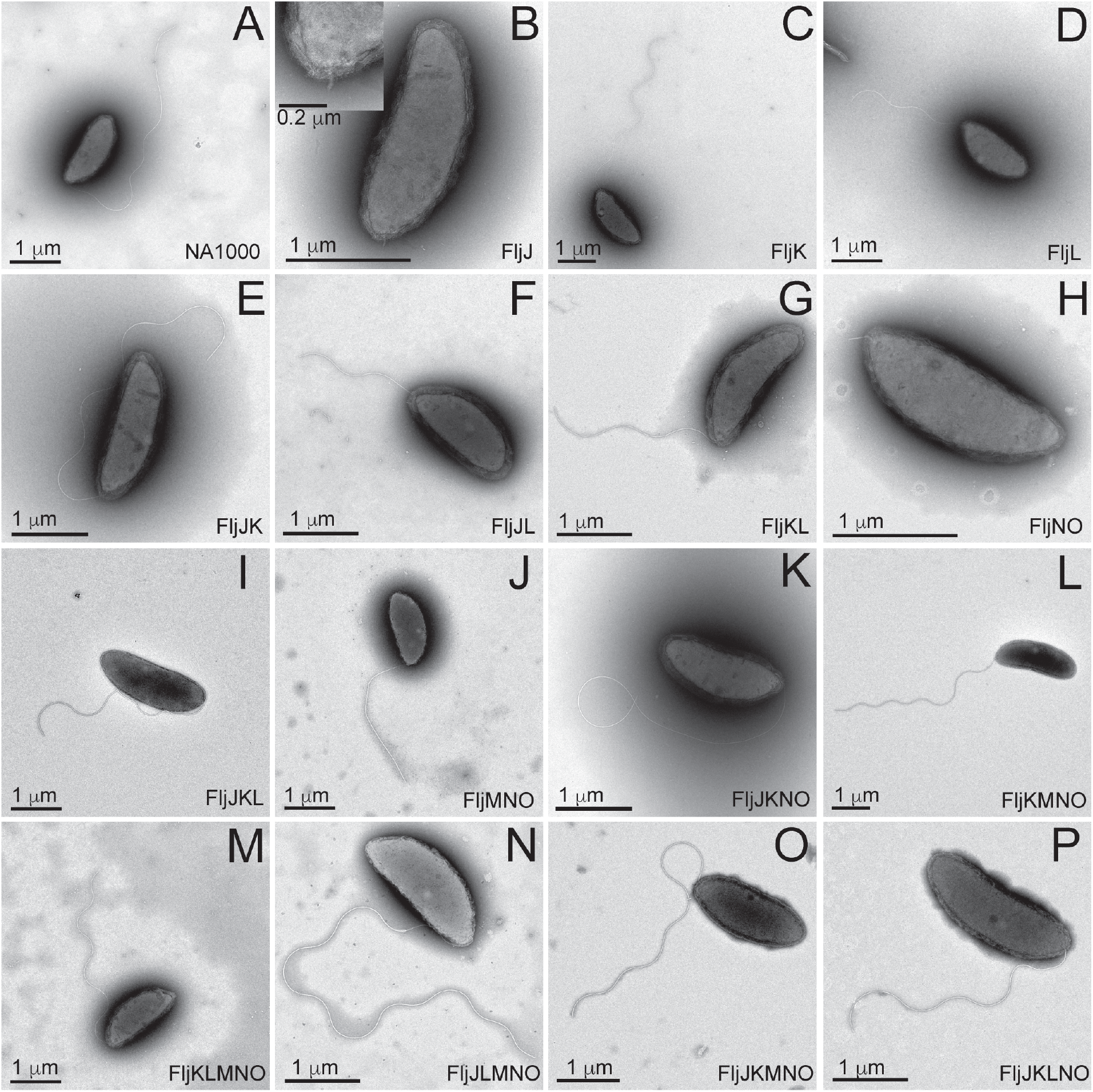
Transmission electron microscopy images of negatively stained *C. crescentus* strains from Figure 1 using 1% phosphotungstic acid, showing that most strains produce flagella. Only FljJ (B) and FljNO (H) presented a severely truncated flagellum that lacked an appreciable flagellar filament. Scale bars are 1 μm unless otherwise indicated.

To further examine the requirements for successful phage adsorption to the *C. crescentus* flagellum, we measured ϕCbK adsorption to a series of four strains that expressed only two flagellins. The FljJK and FljKL strains exhibited the highest motility, highest swimming speed (37), longest filaments and greatest ϕCbK adsorption. Interestingly, FljJK displayed near wild type levels of ϕCbK adsorption that could be attenuated approximately two-fold by replacement of FljJ with FljL. Phage ϕCbK adsorption was significantly reduced for FljJL (34%) and FljNO (1%), which tracked strongly with reduced length, motility, and speed (37).

Increasing in complexity, two strains were examined that expressed three flagellins, FljJKL and FljMNO, which correspond to the alpha or beta locus flagellin genes, respectively (37, 46). The beta locus flagellin strain FljMNO uniformly exhibited decreased speed, motility, filament length, and ϕCbK adsorption relative to wild type. The phenotype of the FljJKL strain, however, yielded two interesting observations. First, deletion of beta locus flagellins confers a minimal effect on speed, motility, and filament length relative to wild type. Secondly, comparison of FljJKL and FljJK shows that including the FljL flagellin attenuates ϕCbK adsorption two-fold relative to the FljJK strain, despite other properties nearly matching wild type (Figs. 1, S1 and S2).

We then examined ϕCbK adsorption to FljKMNO, which contains only the 25 kDa flagellins (Fig. 1C). Motility of FljKMNO was ~50% that of wild type and ϕCbK adsorption was 65% that of wild type (Figs. 1D, S1 and S2). Simultaneously, we studied the profiles for FljJKNO and determined that even though the strain’s motility was higher, at 85%, the adsorption profile was equivalent to FljKMNO (65%).

To quantitatively determine the relationship between the removal of single flagellins and ϕCbK adsorption, we examined FljKLMNO, FljJLMNO, FljJKMNO, and FljJKLNO. Removal of FljN or FljO as single flagellins was not possible, as previously noted by Faulds-Pain (37). Omission of FljJ reduced motility and adsorption to ~80% and 50% that of wild type, respectively. Omission of FljK caused a nominal effect on filament length and motility, but a significant reduction in phage adsorption. Omission of FljL or FljM exerted only minor effects on function and morphology of flagella.

The impact of the inclusion of FljK in the flagellum to ϕCbK adsorption is especially apparent when we compared the slope values from the ϕCbK adsorption kinetics plots for all four strains expressing five flagellins and wild type *C. crescentus* (Fig. S2). The slope for infected wild type cells is −0.0142, the slope for the four strains; in increasing slope steepness are FljJLMNO (−0.0053), FljKLMNO (−0.0119), and FljJKLNO (−0.0183), FljJKMNO (−0.0189). These data illustrate that ϕCbK adsorption is generally lower in the presence of FljL and higher in the presence of FljK, with the noted exception of wild type.

The results of Driks *et al*. indicate that the intermediate size (27 kDa) flagellin (FljL) may serve as an adapter between FljJ and the subsequent assembly of the smaller (25 kDa) flagellin subunits (Fig. 1C). However, our imaging results demonstrate that mutants with deletions of either FljJ or FljL produce flagellar filaments of lengths comparable to that of wild type (Figs. 1C and 2). As such, inclusion of FljJ or FljL is not essential for filament formation despite their localization at the base of the assembled wild type filament.

### Tomographic reconstruction of *C. crescentus* strains during ϕCbK adsorption

The adsorption of ϕCbK phage along the flagellar filament of the flagellin deletion strains was examined by cryo-ET. We resolved intact flagella for all tested deletion strains except for FljJ (Fig. 2B) and FljNO, which had flagella stubs and no ϕCbK phage adsorbed headfirst along the truncated flagellum length. It was noted that ϕCbK phage were adsorbed along the length of the flagella and that the overall structure of the flagella was comparable to the flagella of wild type cells (Fig. 2). It was also determined that a greater number of ϕCbK phages were adsorbed along the flagellum of the FljJK strain (data not shown) when compared with the FljJLMNO mutant, which correlated with the results from the phage adsorption assays. Cryo-ET imaging of ϕCbK-infected strain FljJLMNO revealed that the head filament of ϕCbK could still interact with the flagellum in the absence of FljK, albeit at lower abundance, as expected from our adsorption assays.

Driks *et al*. (47) proposed that the largest (29 kDa) flagellin subunit (FljJ) is used to initiate the assembly of flagellin subunits in *C. crescentus* by forming a short flagellar segment to which other flagellins assemble (Fig. 1C). Our tomographic reconstructions of the FljJ mutant comport with this model, and we resolved the combined length of the corresponding flagellar filament and hook to be between 120 nm and 160 nm, respectively (Fig. 3B and Supplementary Movie 1). Interestingly, the FljJ mutant supports ϕCbK adsorption, albeit at a greatly reduced level when compared with the other strains (Figs. 1C, S1 and S2). The low level ϕCbK adsorption is likely due to ϕCbK tail association with the *C. crescentus* pilus filament and the retraction of the pilus towards the cell pole and the pilus portal, which is the site of irreversible ϕCbK attachment (30, 60, 61).

**Figure 3.**
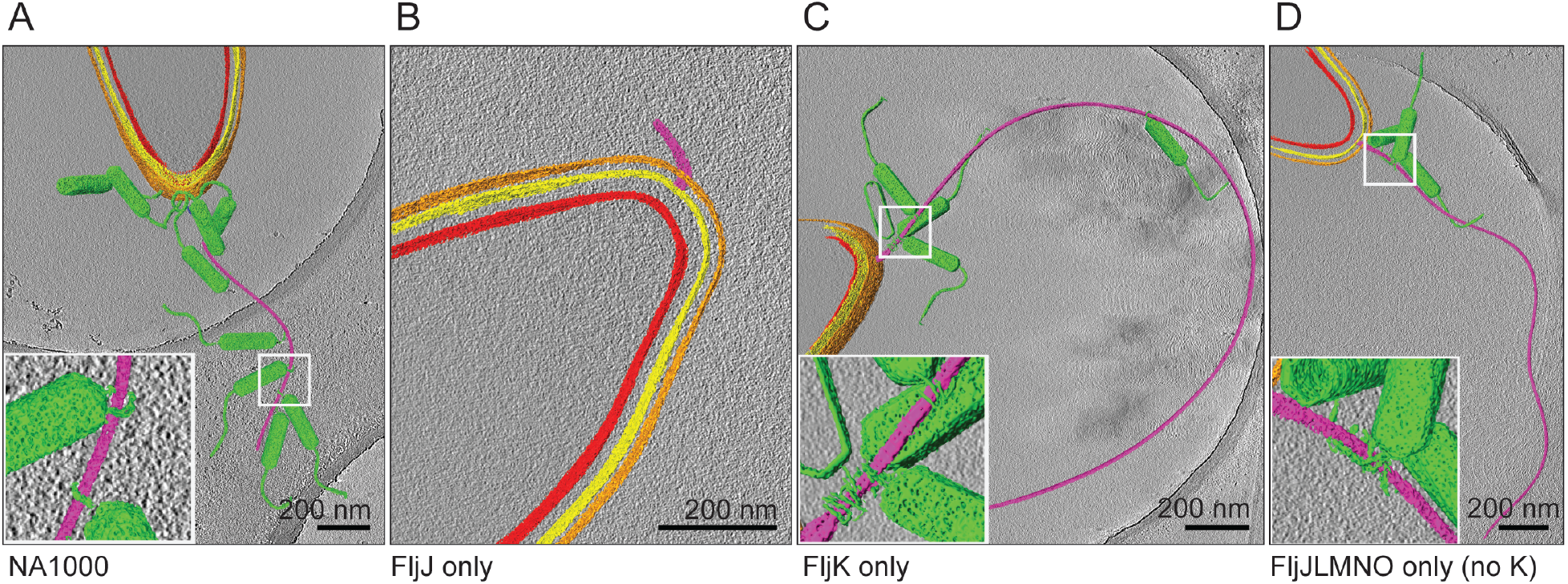
Averaged slices merged with segmentations through three-dimensional tomographic reconstructions of representative ϕCbK-infected *C. crescentus* cells, selected from all strains examined. Segmentation was performed in Amira, coloring the inner membrane red, outer membrane yellow, S-layer orange, flagellum purple, and phage particles green. A, C and D) Phages were observed either adsorbed along the length of the flagellum or attached to the cell poles of strains investigated. B) *C. crescentus* FljJ assembles a short flagellum. Insets highlight the ϕCbK head filament wrapped around the *C. crescentus* flagellum. Scale bars are 200 nm.

### Helical reconstruction of a *C. crescentus* filament containing only FljK

The above data suggest that FljJK is an optimal, but not exclusive, substrate for recruitment of ϕCbK. This is supported by the observation that it is possible to recover approximately 100% of adsorption with a minimal complement of flagellins, so long as at least one is FljK (see stain JK, Fig. 1D,F). We therefore sought to determine a high-resolution structure of FljK in order to build a platform for subsequent interrogation of the filament-ϕCbK interaction mechanism.

Single particle helical reconstruction methods are a powerful tool for high-resolution structure determination of filaments, but most successfully determined structures of homologous flagellins to date have employed straightening mutations to facilitate computational aspects of the reconstruction process (9, 55, 56). We therefore utilized error-prone PCR to mutagenize a plasmid (pCS1) encoding a T103C variant of FljK that is amenable to fluorescence and cryo-correlative light and electron microscopy (cryo-CLEM) imaging via maleimide dye-labeling of the cysteine residues (data not shown) (62). Several isolates were defective for motility and both fluorescence and negative stain TEM imaging were used to determine if such strains indeed harbored straightened filaments (data not shown). One such strain, FljK T103C N130S exhibited straightened filaments which were used for high-resolution cryo-TEM imaging (Fig. 4A).

**Figure 4.**
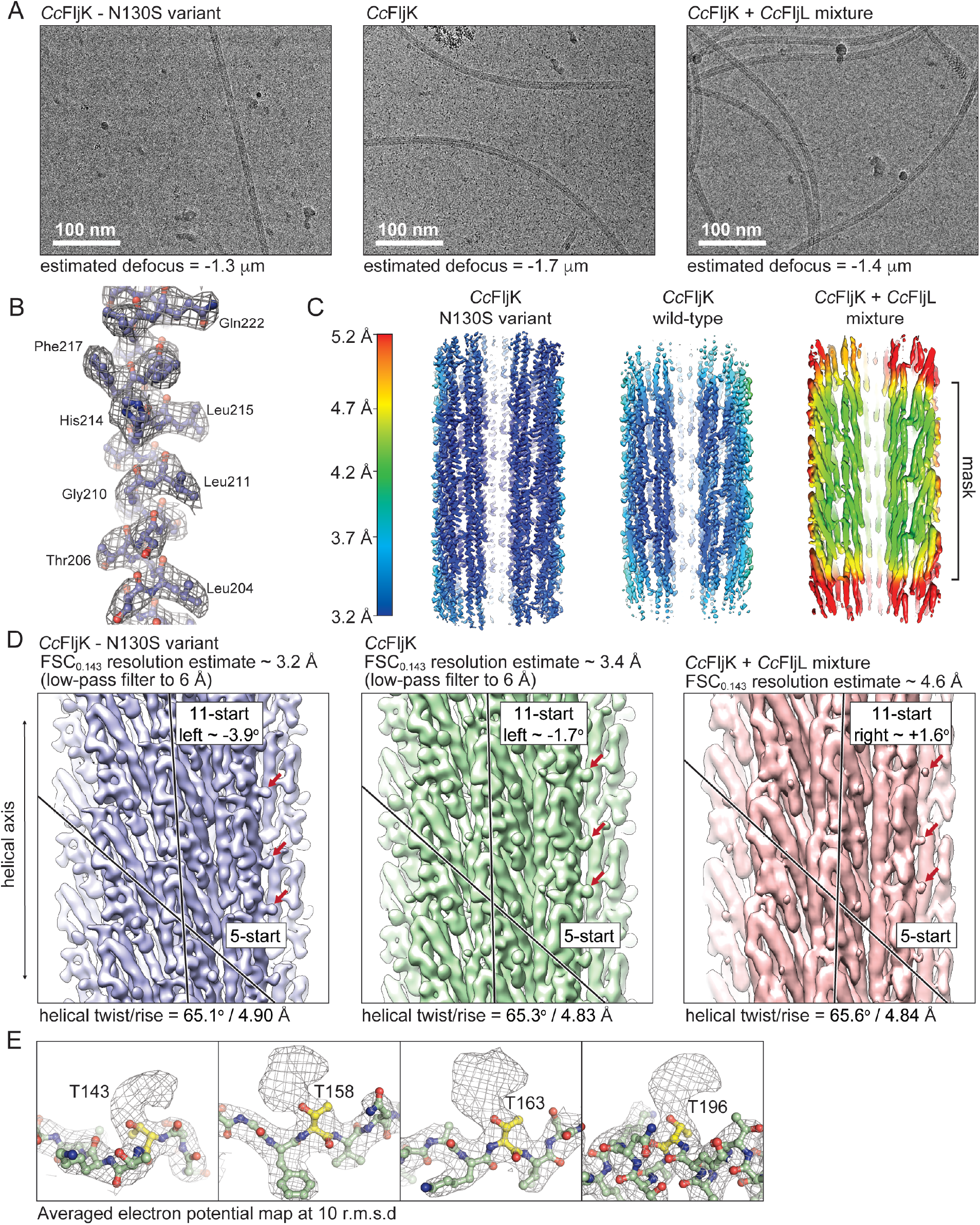
High-resolution imaging and reconstruction of the FljK flagellin. A) Exemplar micrographs of straightened FljK, wild type FljK and a heterogeneous mixture of FljK and FljL. B) example of electron potential map in which main-chain connectivity and side chains are clearly visible. C) Local resolution plots of the three reconstructions generated in Relion 3.1 and Chimera (73, 79). The straightened FljK flagellin exhibits the highest resolution. The heterogeneous mixture of FljK and FljL does not yield a high enough resolution map for modeling of amino acid side chains. D) Comparison of the three reconstructions generated in this work. The single component FljK reconstructions are low-pass filtered to allow visualization of 5- and 11-start protofilaments that are a hallmark of related flagellins (9, 54–56). Note that the heterogeneous FljK/L reconstruction lacks definition of amino acid side chains even in the absence of low-pass filtering. Interestingly, however, the FljK/L reconstruction shows nodules (denote by red arrows) that correspond to apparent glycosylation loci. E) Numerous post-translational modifications are clearly visible on surface exposed threonine residues that are likely O-linked glycans in the FljK only reconstructions.

In lieu of direct helical indexing, we used initial estimates of helical symmetry obtained from high resolution structures of homologous flagellins (9, 54–56) and an initial featureless cylindrical model (Fig. S3). After reconstruction and refinement of helical symmetry parameters in Relion, we were able to recover a map that contained clear definition of amino acid side-chains (Fig. 4B,C), which was utilized as a cross-validation metric for the inherently biased approach used here for acquisition of the initial helical symmetry parameters. We note that the final helical symmetry obtained here is much closer to that of hydrated homologous flagellins (rise/twist ~4.8 Å/65°) than that of dehydrated *C. crescentus* flagellins in negative stain TEM (rise/twist ~3.8 Å/65°) (45).

The straightened N130S variant of FljK exhibited a “locked” left-handed 11-start helix with an approximate inter-subunit rotation of −3.9° (Fig. 4D). Subsequent to reconstruction of this straightened mutant of FljK, we attempted helical reconstruction of a filament that was not “straightened” by an amino acid substitution and therefore not presumably “locked” into either a right- or left-handed 11-start conformer. The resulting reconstruction was of slightly lower resolution and overall map quality relative to the straightened variant of FljK, which is likely due to conformational heterogeneity that is present in the non-straightened filament. However, it was still possible to clearly delineate amino acid side-chains in the majority of the reconstructed map. Interestingly, the wild type variant of FljK exhibited a left-handed 11-start helix with an approximate rotation of −1.7°. The resulting structures of the FljK flagellin monomers and assembled filaments conform to the known architecture of other bacterial flagellins and flagellar filaments (9, 54–56).

We note the appearance of numerous apparent post-translational modifications that decorate the periphery of the FljK filament (Fig. 4E). These modifications are all localized to conserved threonine residues and are likely O-linked glycans, as observed in other flagellin reconstructions (22–24). As the periphery of the filament has the poorest map quality, we were unable to identify the exact glycans nor model them into the atomic models generated here. Future biochemical work will be required to clarify the exact identity of these post-translational modifications.

### Helical reconstruction of a compositionally heterogeneous filament

From our functional data, we find that inclusion of FljL generally impedes adsorption of phage ϕCbK (Fig. 1). For example, comparison of the FljJKL and FljJK strains shows that inclusion of FljL reduces phage ϕCbK adsorption approximately two-fold while exerting a small improvement in motility. We therefore sought to understand what differences may exist at the structural level when FljL is expressed with FljK. Using the same processing pipeline as above, we obtained a helical reconstruction of FljKL flagella that is limited to a final resolution of 4.6 Å. The reduced resolution is attributed to the confounding effects of compositional and conformational heterogeneity, and the reconstruction should be interpreted with caution as it was not indexed *de novo*, nor does the final map exhibit side-chain densities to serve as cross-validation for the reconstruction pipeline used here. However, the low-resolution map still provides some interesting observations. First, the flagellin fold, with terminal D0 domains and intervening D1 domains, is apparent in the reconstruction, noting that the starting model of a featureless cylinder contained no such information. As such, inclusion of FljL does not substantially alter the overall fold or arrangement of monomers in the resulting filament. Second, there are clear nodules on the periphery of the lower-resolution K/L map that map to the same locations as the apparent post-translational modifications in the FljK reconstructions (Fig. 4E). Third, the mixed K/L filament exhibits an 11-start protofilament with a rotation of +1.6°, which is distinct from those of the compositionally-homogeneous FljK reconstructions (Fig. 4D).

### Interprotein contacts in the FljK flagellum

Comparison of the straightened and non-straightened reconstructions of FljK shows that the N130S substitution exerts a minimal effect on the overall structure of FljK, with an all-atom r.m.s.d. of 0.7 Å between the two chains (Fig. 5A). However, there are local differences at protein-protein interfaces that alter the overall organization of the filament. Substitution of N130 for serine disrupts a hydrogen bonding contact between the side-chain of N130 and the carbonyl oxygen of K126 (Fig. 5B,C). This, in turn, disrupts a series of hydrogen bonds between adjacent strands in the 5-start protofilament, resulting in a vertical displacement of approximately 2.5 Å along the helical axis. As such, protein-protein contacts along the exterior of the filament can induce structural changes that propagate to the rest of the helical assembly and change the rotation of the 11-start protofilament. This is noteworthy because phage ϕCbK interacts with the periphery of the filament and could potentially alter the structure of the filament after initial association.

**Figure 5.**
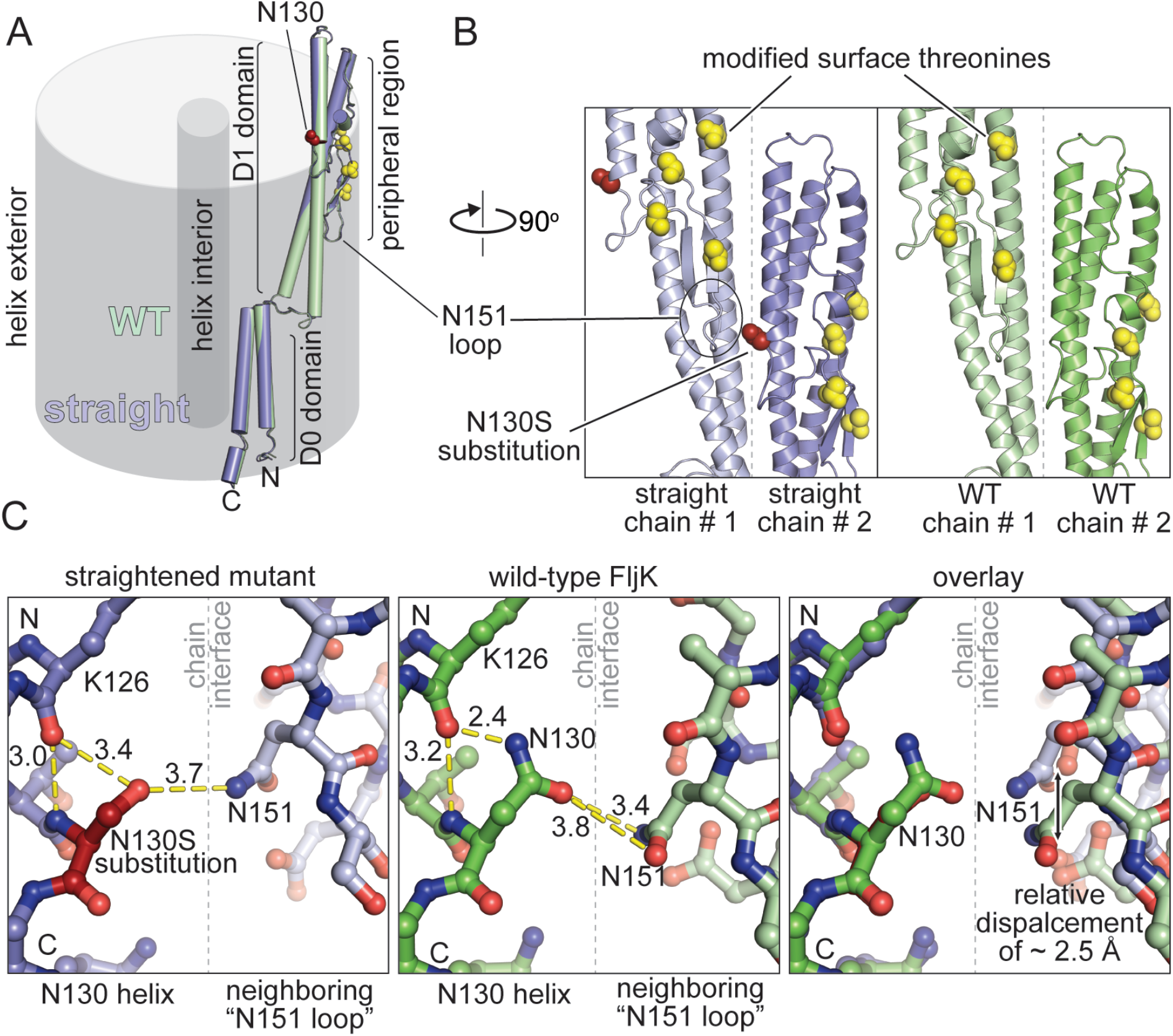
Comparison of local structure in straightened and non-straightened FljK filaments. A) The two FljK variants are highly similar with an all-atom r.m.s.d. of 0.7 Å. The N130S straightening substitution is depicted by red spheres, and post-translationally modified surface threonine residues are depicted by yellow spheres. B) The N130S straightening substitution is located at a protein:protein interface along the surface of the FljK flilament. C) The N130S straightening substitution perturbs a main-chain hydrogen bond with the alpha-helical carbonyl oxygen of K126 and alters the local hydrogen bonding network at a protein:protein interface, resulting in displacement of the neighboring “N151 loop” by approximately 2.5 Å.

## DISCUSSION

### Assembly of a multi-flagellin filament

We have shown that flagellar filament formation in *C. crescentus* is remarkably tolerant of deletions to specific flagellins. For example, all strains imaged here exhibited flagella, with the exception of the FljJ and FljNO strains (Figs. 1 and 2). As such, it appears that alternate arrangements may be possible for filament assembly. Filament assembly in *C. crescentus* is therefore likely a dynamic process with respect to flagellin incorporation, with selection criteria for flagellin insertion that are neither inflexible nor well understood.

We also note a trend upon which inclusion of more types of flagellins generally increases “function” of the *C. crescentus* filament with respect to length, swimming speed, and motility (Fig. 1). The absence of “all or nothing” observations with respect to these properties suggests that the *C. crescentus* flagellum is an exquisitely complex biophysical apparatus that can exploit compositional heterogeneity to enhance function.

### Flagellin complement significantly affects ϕCbK adsorption

Another possible advantage of preserving multiple flagellin paralogs in a bacterial genome is for evasion of flagellotropic phages. For example, a duplicate flagellin may acquire adaptations that attenuate attachment of phage to flagellins, effectively blocking or inhibiting one step of the adsorption process. The alpha locus flagellin FljL appears to play such a role. This viewpoint is supported by two observations. First, a minimal flagellum containing only FljJ and FljK exhibits wild type levels of adsorption, and secondly, addition of FljL to this minimal flagellum attenuates adsorption approximately two-fold while simultaneously increasing flagellum function (as evinced by length and motility). In light of this observation, we have focused our initial high-resolution structure determination efforts on filaments containing FljK and FljL, due to their opposing effects on motility and absorption.

At this point, the role for the beta locus flagellins (FljM, FljN, FljO) remains unclear, as our data exhibit no clear pattern regarding the function of these flagellins. It is possible that the importance of these flagellins may lie in a function not tested here, for example functional redundancy, predation by other bacteriophages, or motility in environments that are not captured by our assay conditions.

### Comparison to other flagellins of known structure

Comparison of our FljK reconstruction to that of other 11-start bacterial flagellins provides several interesting observations. First, helical symmetry is remarkably well-conserved with an average variance in rise and twist below 10% (Fig. 6A). These small differences in helical symmetry, however, can exert a large effect in the handedness of the 11-start protofilament. Until recently, all determined structures of 11-start flagellins fell into two classes: left-handed structures with an 11-start rotation of approximately −1.7° or right-handed structures with an 11-start rotation of approximately +3.8° (9, 54–56). These two dominant classes of 11-start protofilaments strongly influenced the field’s view of filament structure, where toggling between two defined states was thought to play a functional role in bacterial motility (2). Recently, a high-resolution reconstruction of the flagellar filament from *Kurthia sp*. found the corresponding 11-start protofilament to be nearly parallel to the helical axis with a rotation of approximately +0.5°, which could be considered an outlier from all other successful helical reconstructions to date (9, 54–56). Here, we show that two of our reconstructions also fail to conform to the −1.7° or +3.8° precedent for 11-start rotations, and as such there is a previously unappreciated degree of large-scale conformations that flagellar filaments can accommodate, perhaps representing untapped structural diversity in flagellar filament organization (Fig. 6A).

**Figure 6.**
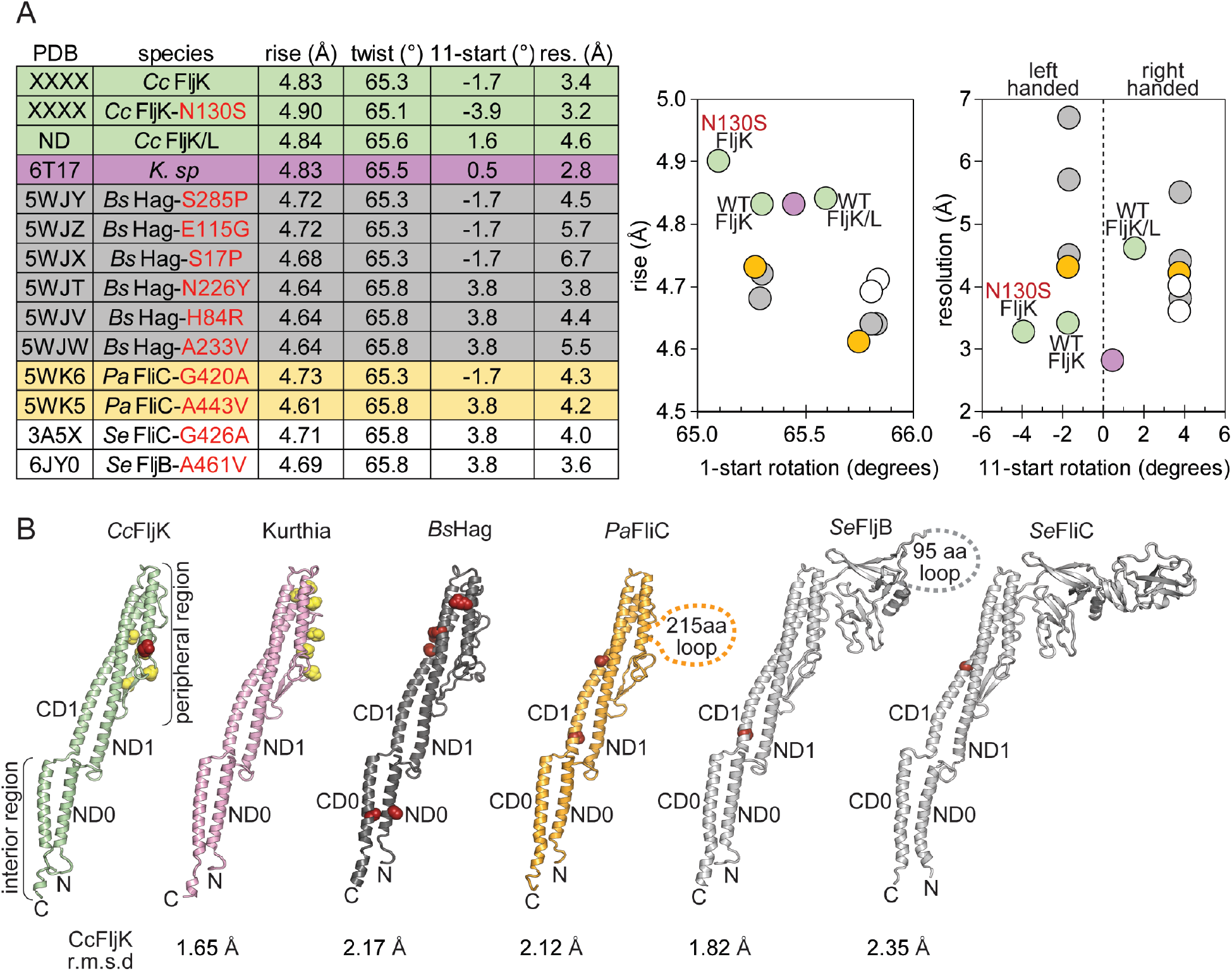
Comparison to other 11-start bacterial flagellins. A) All 11-start bacterial flagellins of known structure have similar helical symmetry values. Neighboring plots are constructed to highlight differences in symmetry. B) Structural comparison of *C. crescentus* FljK to other bacterial flagellins. Known straightening substitutions are depicted by red spheres and post-translationally modified residues are depicted by yellow spheres. The N130S straightening substitution in FljK is a relative outlier in that it perturbs a protein-protein contact at the surface of the flagellin, while the majority of other straightening substitutions typically alter protein-protein contacts in the core of the flagellin.

Despite these differences, we find that flagellins are remarkably well-conserved at the level of secondary structure within the interior of the filament (Fig. 6B). For example, the much larger D2/D3 domains that are present in *Pseudomonas* and *Salmonella* flagellins do not significantly alter the structure of the D0 and D1 regions, which are quite similar to those in the *Caulobacter, Kurthia*, and *Bacillus* flagellins (Fig. 6B). However, the overall architecture of the filament is clearly not insensitive to alterations in the peripheral regions, as our “straightening” N130S substitution is located on the surface of the filament (Fig. 6B). Furthermore, helical assemblies can be so exquisitely sensitive to small changes in local structure that it is possible to completely rearrange the entire helical superstructure through substitution of only a single amino acid (63). We therefore propose the surface threonine-linked modifications play a functional role in altering the overall architecture of the filament. The role of threonine-linked modifications with respect to phage adsorption and interaction is also an interesting topic for future research.

### Helical reconstruction in the absence of *de novo* indexing

Helical reconstruction relies upon *de novo* indexing and acquisition of helical symmetry parameters, a process that is labor intensive and not always readily achievable (51). As such, the current state of helical reconstruction methods very much resembles that of macromolecular X-ray crystallography several decades ago, specifically the requirement for experimental phase determination (64). Here we demonstrate that utilizing prior knowns from homologous structures can greatly accelerate the path to structure determination. More specifically, we have employed something akin to “helical molecular replacement” to obtain initial estimates for helical symmetry from homologous structures to kick-start our reconstructions (65, 66). This approach to helical reconstruction is similarly fraught with biases that can potentially yield false “solutions” for structure determination (50). As such, cross-validation metrics are essential to the pipeline we have employed here. The primary metric we have relied upon is the appearance of high-frequency structural features in our final reconstructions that comport with known stereochemistry and other features of protein structure (Fig. 4B). Furthermore, we also confirmed that observed amplitude spectra from 2-D classification before and after application of helical reconstruction are qualitatively similar (Fig. S4), and as such the application of helical symmetry does not introduce a bias that fundamentally alters the data.

Our ability to generate a reconstruction of a non-straightened FljK flagellum shows that conformational heterogeneity does not necessarily preclude structure determination of flagellar filaments by helical reconstruction methods, similar to recent work by Blum *et al*. (54). We note that map quality is globally degraded in the presence of conformational heterogeneity, and the “best” region of the map becomes more strongly restricted to the center of the reconstruction as conformational heterogeneity increases (Fig. 4C). Excessive conformational heterogeneity can make helical reconstruction impossible, and in such cases conventional single particle reconstructions must instead be utilized (67). However, the latter approach is typically only feasible for larger asymmetric units and, as expected, our attempts to use standard single particle methods on our filaments failed (data not shown) (50).

In comparing our three reconstructions, it is apparent that the combination of compositional and conformational heterogeneity significantly impedes high resolution structure determination. For example, the non-straightened FljK and FljKL filaments exhibit similar curvature (Fig. 4A), yet only the compositionally homogeneous FljK filament yielded a three-dimensional map of sufficient quality to generate an atomic model. One possibility for the limitation of FljKL could be that the highly similar, yet distinct, flagellin monomers are stochastically incorporated throughout the body of the filament, and as such the resulting numerous combinatorial possibilities preclude significant accumulation of any one single composition for averaging *in silico*.

### Sequence analysis of *Caulobacter* flagellins

Our high-resolution reconstructions of FljK allow us to perform a more informed sequence analysis of *C. crescentus* flagellins than was previously possible (Fig. 7A). Interestingly, the surface exposed threonine residues that harbor post-translational modifications in our reconstruction of FljK (Fig. 4E) are conserved in the beta locus flagellins (FljM, FljN, and FljO) and also FljL, but are almost entirely absent in the most divergent flagellin, FljJ. Because nearly all flagellins may be modified, this suggests that the mechanism by which FljK enhances phage ϕCbK predation may not occur through selective recognition of this modification. However, it remains to be seen if phage ϕCbK is sensitive to the threonine linked post-translational modifications that decorate the periphery of the filament.

**Figure 7.**
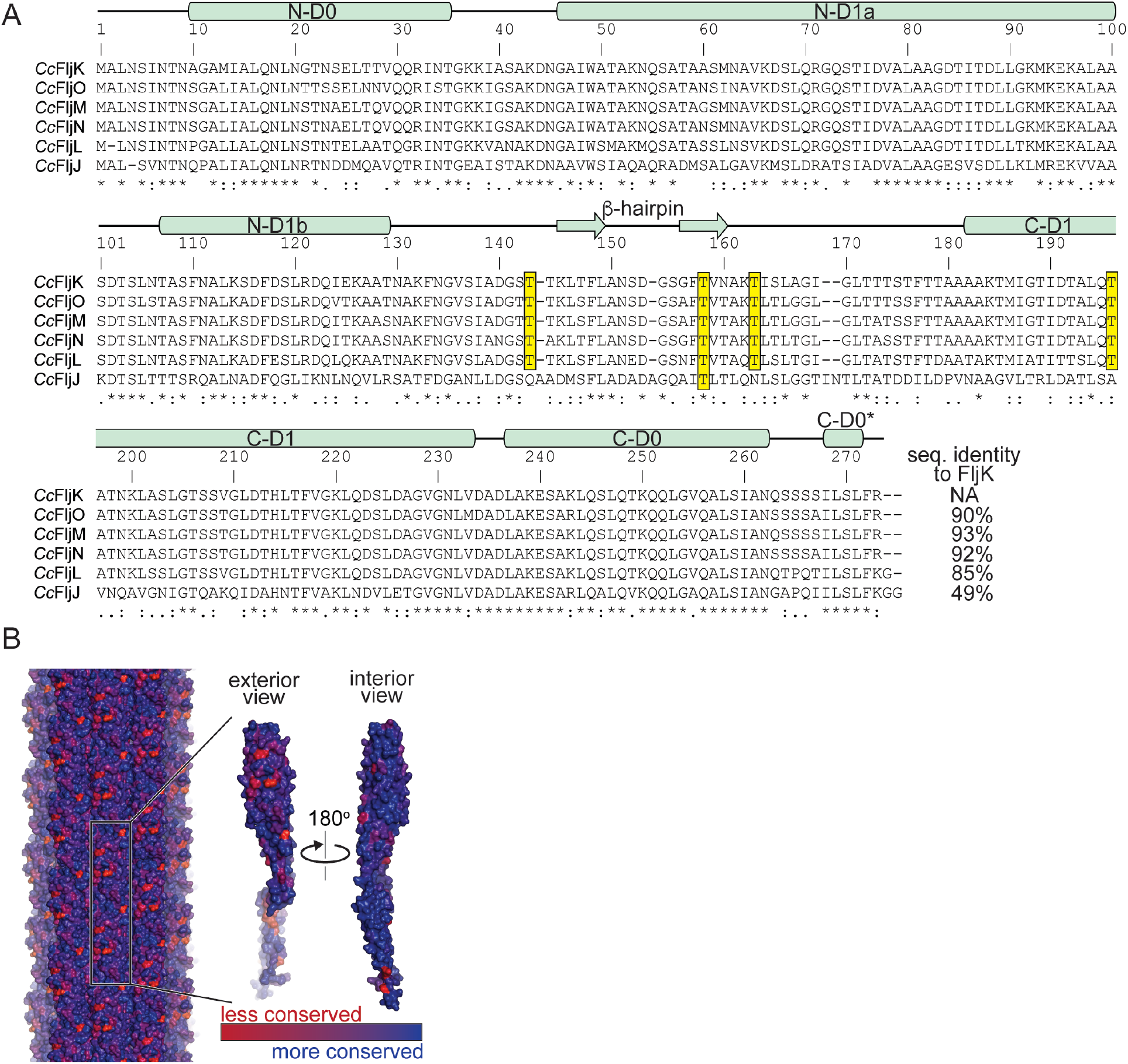
Phylogenetic analysis of flagellin paralogs in *C. crescentus*. A) Sequence alignment of the six flagellins present in *C. crescentus*. The observed secondary structure in FljK is annotated above the sequence alignment. Conserved surface-exposed threonine residues that harbor post-translational modifications are highlighted yellow. Note that the most divergent flagellin, FljJ, lacks most of such threonine residues. B) Conservation mapping of *C. crescentus* flagellins with the program ConSurf (62). The greatest degree of sequence divergence is located along the surface of the flagellin.

We mapped sequence conservation amongst the six *C. crescentus* flagellins onto our structure of FljK and found that the periphery of the filament contains several “hotspots” for sequence variation relative to the interior of the filament (Fig. 7B). As such, differential inclusion of flagellins can alter the surface that is presented to phage ϕCbK. These “hotspots” are an attractive location for future work that seeks to determine the preference of some flagellins by ϕCbK. However, ϕCbK is not absolutely dependent upon any one such surface because the vast majority of strains examined here still exhibit some degree of adsorption. We therefore propose that ϕCbK is somewhat tolerant of variations to the flagellins present on the surface of the *C. crescentus* flagellum.

We also note that FljK and the beta locus flagellins all have N- and C-termini that are conserved in length. This is not unexpected as both termini of these proteins are buried in the lumen of the flagellum. However, comparison of these flagellins with those expected to be located at the base of the filament, FljJ and FljL (Fig. 1C), shows that their amino termini are shorter and carboxy termini longer (Fig. 7A). We speculate that these differences exist for the purpose of altering the local curvature and flexibility of the filament close to the hook and membrane embedded motor, and therefore likely play a role in the transmission of force to the body of the filament.

### Concluding remarks

This work provides numerous insights into determinants of ϕCbK infection of *C. crescentus*, but also raises a host of new questions. For example, what drives ϕCbK preference for FljK and the negative effects of FljL? What is the exact identity of the post-translational modification and is it uniformly or differentially installed along the body of the flagellar filament? What are the functional consequences of these modifications? It remains to be seen if addition of these post-translational modifications can have an effect on motility as observed in *C. difficile* (68), or if they are required for phage association and filament assembly like in *C. jejuni* (69, 70).

We have established a tractable platform for the isolation of various flagellar filaments from *C. crescentus* and rapid structure determination via cryo-electron microscopy. We found that both conformational and compositional heterogeneity are potential roadblocks to overcome when determining helical structures. This platform will allow future biochemical studies to discern the identity of flagellar post-translational modifications in *C. crescentus* and interrogate their functional importance. It will also enable the study of interactions between phages and the flagellar filament, in addition to allowing for the determination of other single flagellin filament structures that do not promote phage ϕCbK adsorption to the same degree as FljK. This information will be valuable to help discern the nuanced, yet functionally relevant, determinants for flagellotropic phage predation in bacteria with compositionally heterogeneous flagella.

## MATERIALS AND METHODS

### Bacterial strains and growth conditions

All *C. crescentus* strains were grown in peptone yeast extract (PYE; 0.2% peptone, 0.1% yeast extract, 1 mM MgSO_4_, and 0.5 mM CaCl_2_) modified from Schmidt and Stanier (21, 55). *E. coli* strains were grown on LB medium (BD, Franklin Lakes, NJ). Selection with Kanamycin, 5 μg/mL for *C. crescentus* and 50 μg/ml for *E. coli*, was applied where indicated. Phage adsorption assays were performed in PYE containing 4 mM instead of 1 mM of MgSO_4_. Plating medium contained 1.5% (w/v) Bacto agar.

### Motility assays

To test the motility phenotypes of flagellin mutants, we carried out motility assays as previously described (4, 30). Briefly, PYE soft-agar plates containing 0.3% (w/v) agar were stab inoculated with isolated colonies from plates that had been incubated overnight. Motility zone diameters were measured on three separate plates after 72 hours of incubation at 30°C and are presented as percentages of the motility of wild type strain NA1000 (Fig. 1D).

### Evaluation of adsorption kinetics

Phage adsorption to the *C. crescentus* flagellum was measured as previously described (30, 32, 71) after the systematic removal of one or more flagellins (Fig. 1D). Briefly, overnight cultures of *C. crescentus* were infected with ϕCbK at a multiplicity of infection (MOI) of 0.002 in PYE adsorption media (4 mM MgSO_4_). A parallel control flask contained only phages. Adsorption proceeded with slow shaking (60 rpm) at 30°C for 45 minutes. At five-minute intervals, aliquots of 100 μl were taken and mixed in ice cold tubes containing 900 μL of PYE medium and 30 μL of chloroform to disrupt bacterial cells; after 45 minutes, samples were centrifuged (5 minutes at 12000 x *g*, 4°C) to precipitate bacterial debris. The supernatant containing unabsorbed phages was used to infect an indicator strain (*C. crescentus* bNY30A; OD_600_ = 0.3) and overlaid on PYE agar. Plates were incubated at 30°C overnight for enumeration of plaque-forming units (PFU). The adsorption rate constant (k) was calculated by plotting relative plaque-forming units over time and dividing the slope of the regression line by the viable bacterial count.

### Negative stain electron microscopy

Samples were prepared for negative stain TEM by first glow discharging 400-mesh copper, carbon-coated, formvar grids (EM Sciences, Hatfield, PA) for 40 seconds. Five μl of overnight cultures of *C. crescentus* were applied to the grids and allowed to adsorb for one minute before being washed three times with fresh PYE and stained with 1% phosphotungstic acid (PTA; pH 6.8) for one minute. Negatively stained *C. crescentus* cells were imaged on a JEOL JEM-1400 transmission electron microscope (TEM; JEOL, Ltd., Japan) equipped with a LaB6 filament and operated at an accelerating voltage of 120 kV. To capture the total length of the bacterial flagellum, images at a low magnification of 2,000× were digitally captured on a Gatan Ultrascan US1000 (2k × 2k) CCD camera (Gatan, Pleasanton, CA). Images were imported into ImageJ and flagellar length was quantified using the measure analysis option (23). The average flagellum length for each *C. crescentus* strain was determined from at least 25 (except strain FljJ) flagella that were intact and attached to the cell.

### Cryo-electron tomography

To characterize ϕCbK adsorption to *C. crescentus* strains by cryo-ET, bacterial overnight cultures were grown to an OD_600_ of 0.6, infected at an MOI of 10 and incubated without shaking for 15 minutes at 30°C. Four μL aliquots were plunge frozen onto glow-discharged, 200 mesh, copper Quantifoil grids (Quantifoil, Germany) in liquid ethane using a Vitrobot Mark III (FEI, Hillsboro, Oregon). BSA-treated 10 nm colloidal gold was either previously applied onto the grids and air dried or mixed with the sample for homogeneous gold distribution in the ice. Data collection was carried out using a JEOL JEM-2200FS 200 kV FEG TEM with an in-column Omega energy filter (slit width 20 eV). Images were collected with a Direct Electron, LP DE-20 direct electron detector (Direct Electron, San Diego, CA). Images were acquired at magnifications with an effective pixel size equal or less than 7.6 Å on the specimen. A cumulative electron dose between 120 e^-^/Å^2^ and 140 e^-^/Å^2^ was used for tilt series acquisition from −62° to +62° and images were acquired at −6.0 μm defocus. Tilt series images were automatically collected with 2° angular increments using SerialEM (24). Tomograms were reconstructed from the aligned images using the IMOD tomography package (72). Three-dimensional volumes were segmented manually using Amira (FEI Visualization Sciences Group).

### Recovery of straight flagella isolates by a genetic screen

*C. crescentus* isolates with straightened FljK flagella were recovered in a screen for loss of motility following mutagenesis. Mutagenesis of *fljK* was performed by error prone PCR on a plasmid (pCS1) that contained the gene encoding the FljK T103C variant, a functional amino acid change that enables flagella to be dyed with a cysteine-reactive maleimide dye (62). Mutagenized pCS1 was transformed into *C. crescentus ΔfljJKLMNO* and transformants were selected by growth on PYE supplemented with 5 μg/ml Kanamycin. Individual isolates were transferred to low percent agar plates as described above for a motility assay. Non-motile isolates of *C. crescentus* were recovered. These isolates were further characterized for straight flagellar filaments by negative stain TEM as described above. The plasmids from non-motile isolates with straightened flagella were purified using the NEB monarch DNA mini prep kit (NEB, Ipswich, MA) and sequenced by Sanger sequencing using the primers (5-GGCACGAATTCCGAGCTGACGACCGTTCAG-3’) and (5’-CAGGCTCAGGATCGAGGACGAAGACTGGTTGGC-3’). One of the non-motile isolates carried a single mutation coding for N130S in *fljK*. This isolate was selected for further testing. The plasmid from this isolate (pCS12) was regenerated in the pCS1 background using quick change site-directed mutagenesis with the primers (5’-TTGAACTTGGCGCTCGTCGCGGCCTTTTCG-3’ and 5’-CGAAAAGGCCGCGACGAGCGCCAAGTTCAA-3’). The plasmid pCS12 was transformed into *C. crescentus ΔfljJKLMNO* and the resulting phenotype was confirmed to be similar to the original non-motile isolate.

### Isolation of flagellins for high-resolution structural analysis

Three liters of *C. crescentus* cells were grown to an OD_600_ of 0.6 in PYE. Cells were pelleted at 10,000 × *g* for 15 minutes and the supernatant was collected. Then, the supernatant was centrifuged at 48,000 × *g* for 45 minutes to pellet flagellar filaments. The resulting pellet was gently resuspended by overlaying with three mL of phosphate-buffered saline and incubating overnight at 4°C; harsh mechanical resuspension was avoided to prevent filament damage. The 10,000 × *g* and 48,000 × *g* centrifugation steps were repeated to remove additional debris. Then, the flagella solution was centrifuged at 17,500 × *g* for 15 minutes and the supernatant was collected, followed by a final pelleting spin at 21,000 × *g* for two hours and gentle resuspension in 100 μL of PBS as above.

### Grid Preparation

Four μL aliquots of the purified flagella were plunge frozen onto glow-discharged, 200 mesh, copper Quantifoil grids (Quantifoil Micro Tools GmbH, Germany) or C-flat grids (Protochips, Morrisville, North Carolina) in liquid ethane using a Vitrobot Mark IV (Thermo Fisher, Hillsboro, Oregon). In some cases, BSA-treated 10 nm colloidal gold was mixed with the sample for homogeneous gold distribution in the ice.

### Data collection and processing

Imaging was performed on a ThermoFisher Titan Krios FEG-TEM operated at 300 kV and a Gatan K3 camera. Dose-fractionated micrographs were collected in counting mode over a nominal defocus range of −0.3 to −2.5 μm at a pixel size of 0.87 Å with a total dose of ~60 e^-^/Å^2^ (or 1.33 e^-^/Å^2^/frame) on the specimen and ~1 e^-^/pixel/frame on the camera (Table 1). Data processing was conducted within the Relion 3.1 framework (50, 72, 73). Gain-normalized micrographs were motion corrected with MotionCor2 (74) and initial estimates for whole-micrograph defocus performed in gCTF (75). Segments were manually picked in Relion 3.1 and extracted with a box size of 320 pixels and an interbox distance of 25 Å. 2-D classification was performed on two-fold down-sampled particles (box size of 160 pixels, 1.74 Å pixel size) using 50 classes with an in-plane angular sampling of 0.25° and a mask of 250 Å (Fig. S3). Segments contributing to the best classes were then re-extracted with re-centering.

**Table 1.** Data collection, reconstruction, and model statistics.

| Protein | CcFljK<br>T103C/N130S | CcFljK<br>wild type | CcFljK + L<br>wild type |
| --- | --- | --- | --- |
| PDB ID | XXXX | XXXX | NA |
| EMDB ID | EMD-XXXXX | EMD-XXXXX | EMD-XXXXX |
| **Data collection and** |  |  |  |
| Voltage (kV) | 300 | 300 | 300 |
| Electron exposure (e<sup>-</sup>/Å<sup>2</sup>) | 60 | 60 | 60 |
| Number of frames | 45 | 45 | 45 |
| Nominal defocus range (-μm) | 0.3 - 2.5 | 0.3 - 2.5 | 0.3 - 2.5 |
| Pixel size (Å) | 0.87 | 0.87 | 0.87 |
| Symmetry (beyond helical) | C<sub>1</sub> | C<sub>1</sub> | C<sub>1</sub> |
| Micrographs | 630 | 1,016 | 494 |
| Initial particles | 82,716 | 177,826 | 160,871 |
| Final particles | 59,868 | 77,146 | 54,743 |
| Box size (Å) | 278 | 278 | 278 |
| Box size (pixels) | 320 | 320 | 320 |
| Helical symmetry |  |  |  |
| rise (Å) | 4.90 | 4.83 | 4.88 |
| twist (degrees) | 65.2 | 65.3 | 65.6 |
| 11-start filament hand | Left | Left | Right |
| Helical Z parameter (%) |  |  |  |
| masking | 70 | 70 | 70 |
| refinement | 50 | 50 | 50 |
| Map resolution (Å) | 3.2 | 3.4 | 4.6 |
| FSC threshold | 0.143 | 0.143 | 0.143 |
| Sharpening B factor (Å<sup>2</sup>) | -64 | -66 | -177 |
| **Structure Refinement** |  |  |  |
| Total atoms | 38,760 | 38,820 | NA |
| ligands | 0 | 0 | NA |
| water | 0 | 0 | NA |
| RMS, bonds (Å) | 0.007 | 0.008 | NA |
| RMS, angles (°) | 1.0 | 0.8 | NA |
| Ramachandran favored (%) | 98.5 | 97.0 | NA |
| Ramachandran outliers (%) | 0 | 0 | NA |
| Rotamer outliers (%) | 0.5 | 0.0 | NA |
| Map-model correlation | 0.86 | 0.80 | NA |
| Model B factors |  |  |  |
| minimum | 19.3 | 27.8 | NA |
| mean | 30.6 | 50.4 | NA |
| maximum | 60.1 | 159 | NA |
| MolProbity score | 1.2 | 1.7 | NA |
| Clashscore | 4.7 | 9.2 | NA |

Amplitude spectra of the best 2-D classes were of insufficient quality for *de novo* indexing (Fig. S3B). Therefore, initial estimates for helical symmetry were taken from homologous flagellins of known structure (9, 54–56). An initial model was obtained using 3-D classification in Relion 3.1, starting from a featureless cylinder and a fixed helical rise of 4.8 Å and helical twist of 65.6° (Fig. 3SC,D). The resulting volumes exhibited clear secondary structure features that are consistent with previously determined structures of flagellins. One such volume was then low-pass filtered to 10 Å and used as an input volume for another iteration of 3-D classification in which the helical symmetry was not fixed. A consensus from the best classes was combined and subjected to helical auto-refinement with optimization of helical symmetry parameters, after which point the resulting maps for the FljK only reconstructions were of sufficient quality to unambiguously trace the main-chain and model the side-chains of most residues. Subsequent to auto-refinement and per-particle CTF and motion correction, another round of 3-D classification without alignment was used to further curate the dataset, prior to re-extraction to the native pixel size of 0.87 Å and a final round of masked refinement and post-processing. Subsequent to reconstruction, a final round of 2-D classification without alignment was used to ensure the refined particle stack exhibited 2-D classes with amplitude spectra that were qualitatively similar to amplitude spectra obtained from naïve 2-D classification before imposing any estimates of helical symmetry from homologous structures (9, 54–56)(Fig. S4). Masked FSC curves are shown in Supplementary Figure 4C.

### Model building

Manual model building was first conducted with the straightened variant of FljK as the corresponding map exhibited the best resolution of side-chain densities throughout the asymmetric unit. Model building was performed *de novo* for a monomer in Coot (76), and Phenix (77) was then used for real space refinement with Ramachandran (78) and secondary structure restraints. After refinement to convergence, the refined monomer was manually placed in all positions throughout the helical reconstruction and subjected to a final round of refinement in Phenix. A model of the wild type FljK was built *de novo* as above, but automated refinement in Phenix additionally employed reference model restraints (77) from a poly-alanine variant of the above straightened monomer. The resulting models exhibit excellent stereochemistry and map-model correlation (Table 1). Model building was not performed for the heterogeneous FljK/L reconstruction, as the corresponding map lacked sufficient definition to assign the position of amino acid side-chains, and as such lacked an essential cross-validation metric for the initial helical symmetry estimates that were used during reconstruction.

## ACKNOWLEDGMENTS

The authors are grateful to J. Kollman and E. Lynch (Univ. of Washington), E. Bullitt (Boston Univ.), and E. Egelman (Univ. of Virginia) for valuable advice and feedback on numerous aspects of helical reconstruction methods. The authors thank Y. Brun (U. Montreal) and P. Aldridge (Newcastle U.) for *Caulobacter crescentus* strains and valuable discussions about bacterial flagella and cell motility. The authors acknowledge the Robert P. Apkarian Integrated Electron Microscopy Core of Emory University for microscopy services and support. This work was supported in part by the University of Wisconsin – Madison, the Morgridge Institute for Research, Emory University, Children’s Healthcare of Atlanta, and the Georgia Research Alliance to E.R.W.; public health service grants R01GM104540 and R01GM104540-03S1 to E.R.W. from the NIH, and NSF grant 0923395 to E.R.W. Data collection at Florida State University was made possible by NIH grants S10 OD018142-01, S10 RR025080-01, and U24 GM116788 to Kenneth A. Taylor.

R.S.D., C.W.S., Z.K., R.C.G-F., J.C.S., E.R.W. prepared strains, conducted motility and functional assays and negative stain imaging. R.S.D., Z.K., R.C.G-F., E.R.W prepared samples for cryo-ET and calculated reconstructions. N.T.P., D.P., J.C.S. prepared samples for helical reconstruction. E.J.M. and N.T.P. collected data for helical reconstruction. E.J.M performed helical reconstruction. All authors contributed to data interpretation and manuscript preparation. E.R.W conceived of and supervised the work.

## DATA AVAILABILITY

Maps for the mutant FljK, wild type FljK, and FljK/L reconstructions have been deposited into the Electron Microscopy Data Bank under accessions EMD-XXXXX, EMD-XXXXX, EMD-XXXXX, respectively. Coordinates for the mutant FljK and wild type FljK models have been deposited in the Protein Data Bank under accessions XXXX and XXXX, respectively. Other data supporting the findings of this manuscript are available from the corresponding author upon request. The authors declare no competing financial interests. Correspondence and requests for materials should be addressed to E.R.W.

