## Supplementary figures for "Flagellar structures from the bacterium *Caulobacter crescentus* and implications for phage ϕCbK predation of multi-flagellin bacteria"

**Supplementary figures and supplementary figure legends**

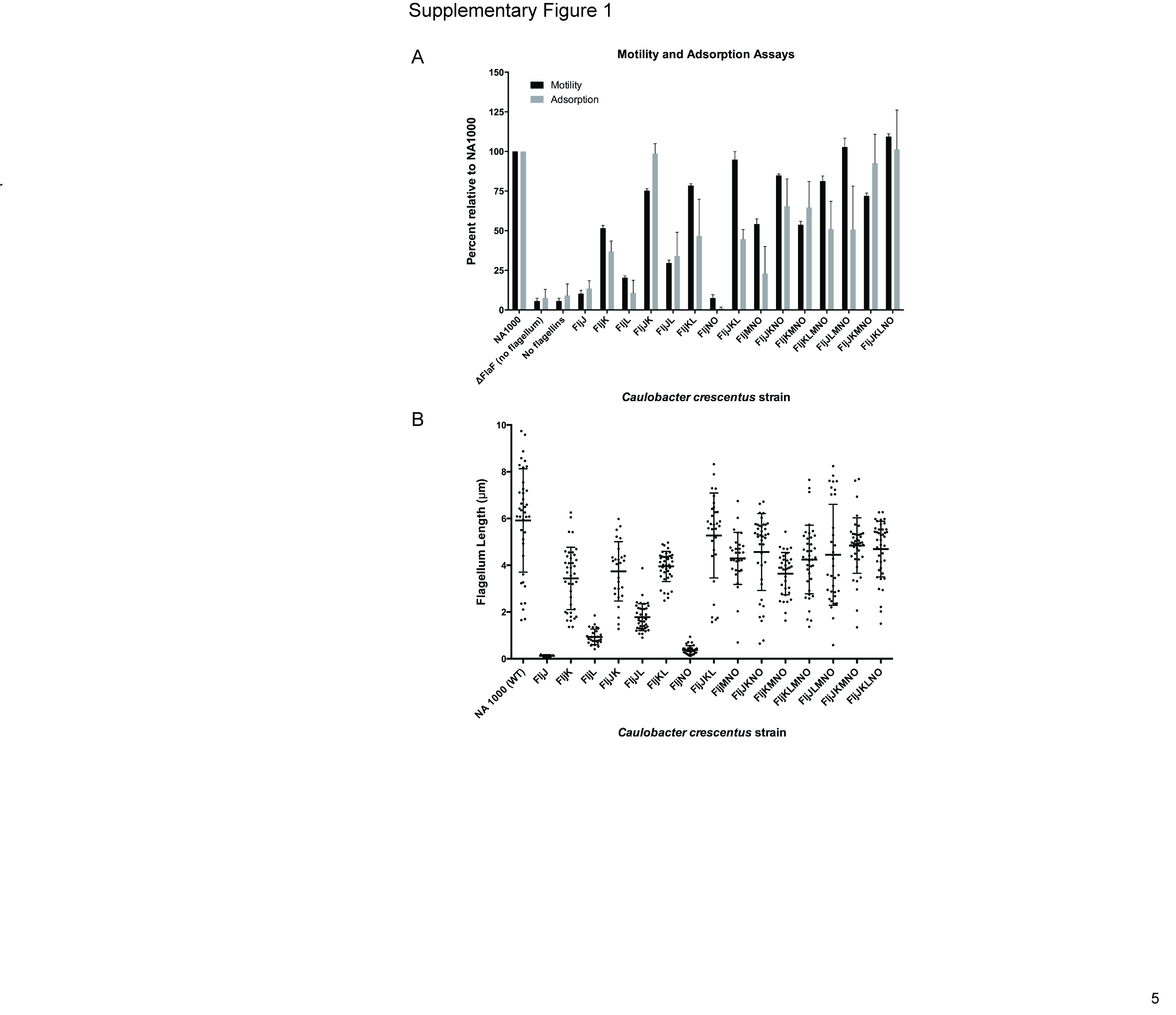

**Supplementary Figure 1.** Cell motility and adsorption kinetics of phage ϕCbK to wild type and flagellin mutants of *C. crescentus*. A) Strain motility (black bars) and adsorption of phage ϕCbK (light grey bars) were measured in triplicate and are normalized relative to wild type *C. crescentus* strain (NA1000). B) Flagellum length measurements relative to wild type NA1000 *C. crescentus*. The black points indicate individual measurements of each flagellum. Error bars indicate standard deviation and the central cross bars represent the average length measurement for each strain. For clarity, the strains are named by the flagellin(s) they synthesize.

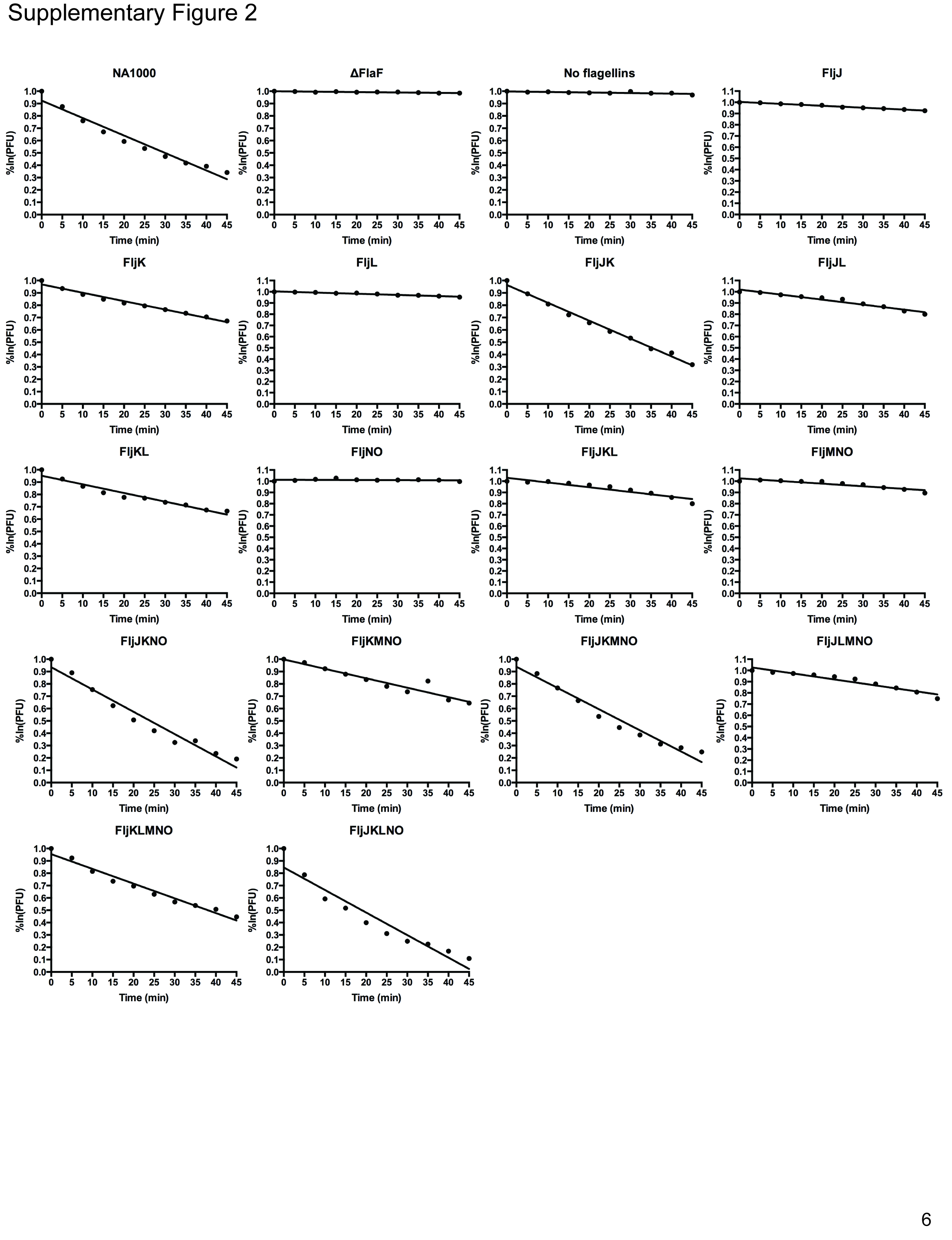

**Supplementary Figure 2.** Adsorption kinetics of phage ϕCbK to *C. crescentus* strains with altered flagellin complements. Data points represent the natural logarithm of phage titer relative to the initial titer at time 0 [ln(T_t_/T_t=0_)].

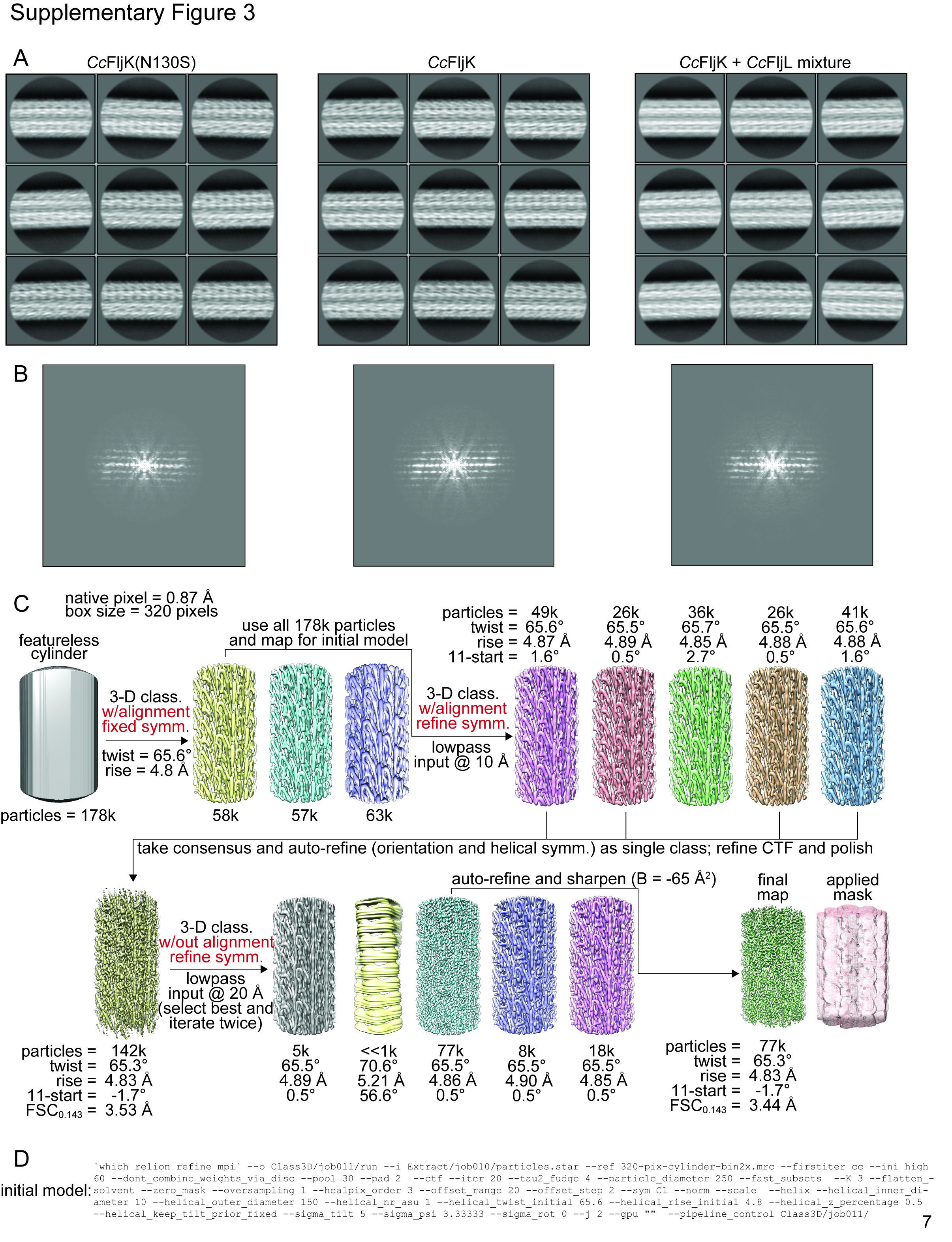

**Supplementary Figure 3.** Helical reconstruction in the Relion 3.1 framework. A) Example 2-D classes. B) Representative amplitude spectra from initial 2-D classes. Although the spectra are not of sufficient quality for *de novo* determination of helical symmetry, the observed layer line spacing is consistent with a helical repeat that is similar to that in homologous flagellins of known structure (1-4)(Supplementary Figure 4). C) Helical reconstruction using initial helical symmetry values from homologous flagellins. After generation of an initial model, the helical symmetry values are further refined to yield a high-resolution structure. Using this pipeline, the FljK-only flagellins give reconstructions in the low 3 Å resolution range, whereas the heterogeneous FljK/L flagellin is trapped at ~ 4.6 Å. D) Example command used for initial model generation in Relion 3.1.

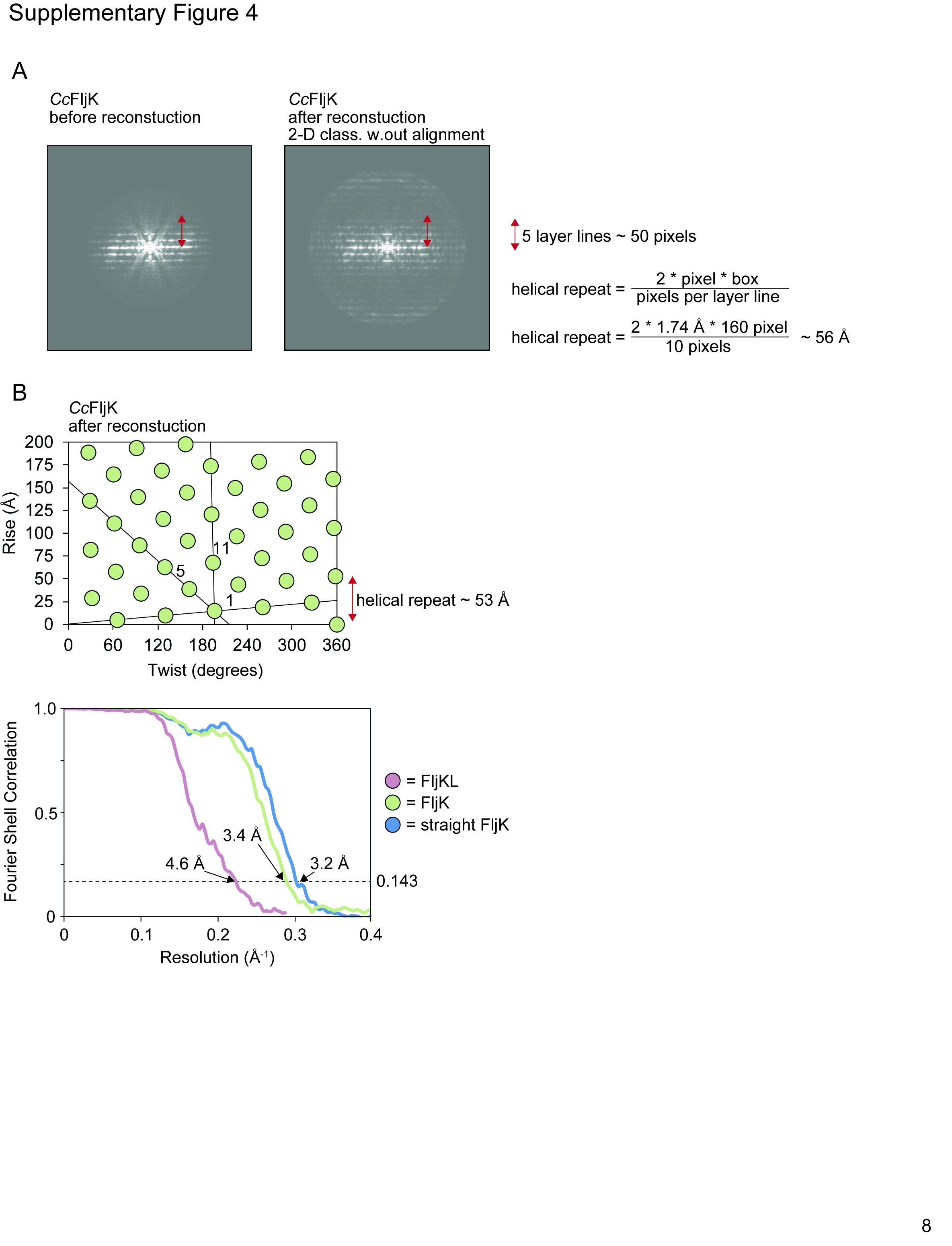
**Supplementary Figure 4.** Comparison of amplitude spectra before and after helical reconstruction. A) Representative amplitude spectrum from a naïve 2-D classification is similar to that obtained from an alignment-free 2-D classification run after helical reconstruction in Relion 3.1. In both cases, the observed repeat distance is ~ 56 Å. B) When viewed in a helical plot, the refined rise and twist from reconstruction gives a similar repeat distance of ~ 53 Å. C) Masked FSC curves for the three reconstructions presented here.

**Supplementary Movie 1**. Three-dimensional tomographic reconstruction and segmentation of the *C. crescentus* FljJ cell presented in Figure 3B. Features are colored to indicate the inner (red) and outer (yellow) membranes, S-layer (orange), and flagellum (fuchsia). Scale bar 200 nm.
